# *HTT* silencing delays onset and slows progression of Huntington’s disease like phenotype: Monitoring with a novel neurovascular biomarker

**DOI:** 10.1101/2020.11.17.386631

**Authors:** Hongshuai Liu, Chuangchuang Zhang, Jiadi Xu, Jing Jin, Liam Cheng, Qian Wu, Zhiliang Wei, Peiying Liu, Hanzhang Lu, Peter C. M. van Zijl, Christopher A. Ross, Jun Hua, Wenzhen Duan

## Abstract

Huntington’s disease (HD) is a dominantly inherited, fatal neurodegenerative disorder caused by a CAG expansion in the *Huntingtin* (*HTT*) gene, coding for pathologic mutant HTT protein (mHTT). Because of its gain-of-function mechanism and monogenic etiology, strategies to lower HTT are being actively investigated as disease-modifying therapies. Most approaches are currently targeted at the manifest HD stage, when clinical outcomes are used to evaluate the effectiveness of therapy. However, as almost 50% of striatal volume has been lost at the time of onset of manifest HD it would be preferable to begin therapy in the premanifest period. An unmet challenge is how to evaluate therapeutic efficacy before the presence of clinical symptoms as outcome measures. To address this, we have been developing more sensitive biomarkers such as functional neuroimaging with the goal of identifying noninvasive biomarkers that provide insight into the best time to introduce HTT-lowering treatment. In this study, we mapped the temporal trajectories of arteriolar cerebral blood volumes (CBVa) using inflow-based vascular-space-occupancy (iVASO) MRI technique in an HD mouse model. Significantly elevated CBVa was evident in premanifest zQ175 HD mice prior to motor deficits and striatal atrophy, recapitulating altered CBVa in human premanifest HD. CRISPR/Cas9-mediated non-allele-specific *HTT* silencing in striatal neurons restored altered CBVa in premanifest zQ175 mice, delayed onset of striatal atrophy, and slowed the progression of motor phenotype and brain pathology. This study showed the potential of CBVa as a noninvasive fMRI biomarker for premanifest HD clinical trials and demonstrates long-term benefits of introducing an HTT lowering treatment in the premanifest HD.

## Main

Huntington’s disease (HD) preferentially involves the basal ganglia- especially the striatum- but also affects other brain regions and has no cure or disease-modifying treatment ^1,2^. While cognitive and emotional changes are increasingly recognized as causes of disability, motor changes remain key and easily quantifiable aspects of clinical onset and progression ^3^. HD is caused by a CAG repeat expansion in the *huntingtin* gene *(HTT)*^2^ and, as a monogenic disorder, can serve as a model for studying other neurodegenerative diseases. Because of the gain-of-function mechanism of mutant HTT, lowering HTT levels is rapidly emerging as a powerful disease-modifying therapy ^4,5^.

Most current therapeutic approaches target the manifest HD stage when clinical outcomes can be used to evaluate the effectiveness of therapy. However, structural magnetic resonance imaging (MRI) studies have shown that almost 50% of striatal volume has already been lost by the onset of manifest HD ^6–10^. Thus, it would be preferable to begin treatment in the premanifest period, and equally important to have biomarkers indicative of treatment response prior to the development of clinical symptoms. Volume of the striatum is a potentially valuable biomarker, but it changes slowly, and the extent to which it will demonstrate response to treatment is unknown. Cerebrospinal fluid (CSF) and blood biomarkers may also be useful ^11–14^; however, their relationship to disease progression and response to treatment remains less well understood. Therefore, we have been developing sensitive and noninvasive biomarkers such as functional neuroimaging ^15^.

The clinical diagnosis of HD is based on the presence of movement disorders. However, functional changes in the brain can precede motor onset by many years ^16–21^. CAG repeat expansion length can be used after predictive genetic testing during the premanifest period to approximate the age of motor onset (manifest HD). Given the availability of predictive genetic testing, mutant *HTT* carriers can be identified decades before clinical manifestation, providing a unique opportunity to identify premanifest biomarkers which can facilitate the development of disease-modifying therapies. Moreover, sensitive biomarkers are crucial for determining the optimal time to begin treatment ^1,17,22–24^.

The brain is a high energy-demanding organ despite its limited intrinsic energy storage ^25^. The balance between substrate delivery from blood flow and neuronal/glial energy demand is precisely regulated in a healthy brain, and cerebral blood flow/volume are strongly coupled with brain metabolism ^26^. A recent study indicated that impaired mitochondrial oxidative phosphorylation in vulnerable striatal neurons occurs long before manifest HD^27^. As arterioles are actively-regulated blood vessels, arteriolar cerebral blood volume (CBVa) may be sensitive to premanifest alterations in the HD brain ^28–30^. We and others have reported elevated CBVa in human premanifest HD ^15,26^.

Several HTT-lowering approaches are or soon will be in clinical trials. These include antisense oligonucleotides (ASOs) and AAV-mediated delivery of RNA interference (RNAi) reagents ^4^ A more recent approach using RNA-guided Clustered Regularly Interspaced Short Palindromic Repeats (CRISPR)/CRISPR-associated (Cas) systems, which induce DNA double-strand breaks and permanently disable mutant gene function in target cells, has been applied in preclinical studies with HD mouse models ^31,32^. It remains an unmet challenge to reliably evaluate the effectiveness of these HTT lowering treatments in clinical trials, particularly in premanifest HD.

HD mouse models have helped elucidate disease pathogenesis, experimental therapeutics, as well as biomarker development ^33,34^. While prior focus has been on neurons in HD, it is becoming increasingly apparent that other components such as glial cells and blood vessels may contribute to or at least be reflective of pathogenesis ^35–37^. The full-length *HTT* knock-in zQ175 mouse model ^38,39^ has the advantage of a relatively slow development of pathology with a period resembling the human HD premanifest stage. We have previously characterized homozygous zQ175 mice and defined progressive motor, neuropathological, and structural neuroimaging changes ^40^. But since most HD patients carry one mutant *HTT* allele, heterozygous zQ175 KI mice more closely resemble the appropriate genomic and protein context of human HD.

In this study, we employed a CRISPR/Cas9 system to lower mutant (and wild type) *HTT* expression in striatal neurons of heterozygous zQ175 mice. We then used inflow-based vascular-space-occupancy (iVASO) MRI (CBVa imaging) in conjunction with behavioral assessments and structural neuroimaging to evaluate the effectiveness of *HTT* silencing introduced at the premanifest stage. We demonstrate that lowering *HTT* in striatal neurons restores CBVa in premanifest zQ175 mice, delays onset, and slows the progression of motor phenotype and striatal atrophy. Our data suggest that CBVa measured by iVASO MRI may be a promising imaging biomarker for premanifest HD clinical trials, and that introducing treatment at the premanifest stage has long-lasting benefits in alleviating manifest HD.

## Results

### Capturing abnormal CBVa trajectories in zQ175 mice

We previously observed elevated CBVa in human premanifest HD brains ^15^. In the present study, we mapped temporal trajectories of CBVa using iVASO MRI measures, and conducted motor phenotype and striatal volume assessments in the heterozygous zQ175 HD model which expresses human *HTT* exon-1 with an expanded CAG repeat. Longitudinal characterization was conducted at 3, 6, 9 months of age, representing prior to onset (premanifest, 3M), onset (6M), and post-onset (manifest, 9M) of striatal atrophy and motor phenotypes (Fig. 1a). Mouse brains were imaged using a 3D acquisition protocol, which allowed continuous scanning of 0.1 mm-thick slices of the whole brain to minimize partial volume sampling effects in the dataset. An ROI-based analysis was performed in the basal ganglion system with focuses on the striatum and motor cortex. We found that CBVa was significantly elevated in the striatum (Fig 1b-c, *p*=0.034) and motor cortex (Fig. 1d-e, *p*=0.041) of zQ175 mice at 3 months compared to wild type littermate controls. No significant differences were observed in the striatal volume, motor function, locomotor activity, and body weight at this age between zQ175 mice and controls (Extended data Fig. 1 a-e). These data indicate that altered CBVa occurs prior to motor symptoms and striatal atrophy in heterozygous zQ175 mice, and that the CBVa changes in this mouse model are reminiscent of those observed in the premanifest human HD brain.

**Figure 1.**
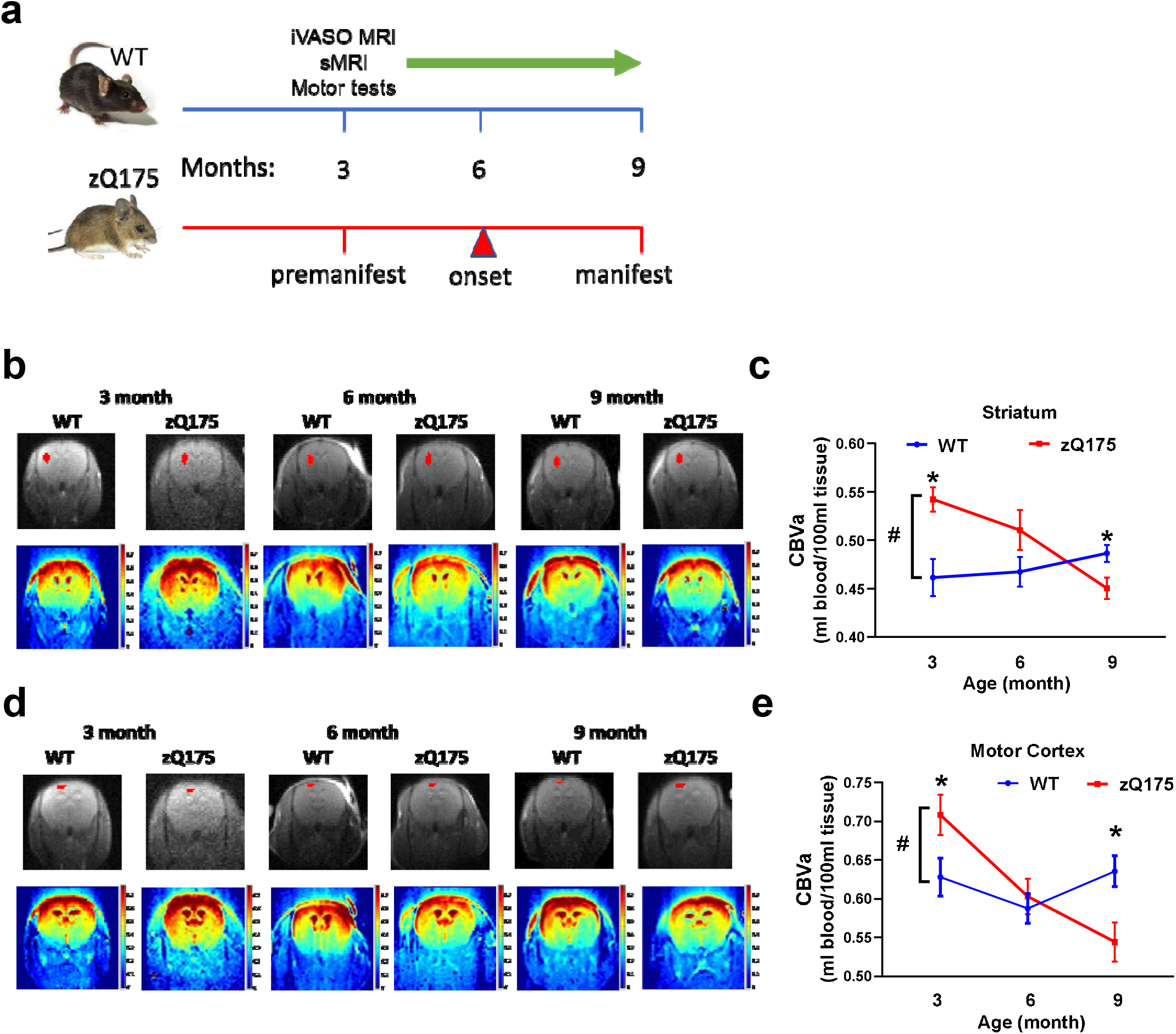
Abnormal trajectories of arteriolar cerebral blood volume (CBVa) in zQ175 mice. (**a**) Timeline for *in vivo* experiments. **(b)** Representative CBVa maps calculated from inflow-based vascular-space-occupancy (iVASO) images in mouse brains from indicated genotypes and ages. Top row shows the raw images, and the red ROIs indicate the quantified brain region-striatum. Bottom row shows the CBVa maps for zQ175 mice and wild type (WT) littermates. The scale bars are shown on the right and warmer colors represent higher CBVa values. (**c)** Longitudinal CBVa changes in the striatum of zQ175 HD mice and WT littermates. (**d**) Representative CBVa maps calculated from iVASO images in mouse brains from indicated genotypes and ages. Top row shows the raw images, and the red ROIs indicate the quantified brain region-motor cortex. Bottom row shows the CBVa maps for zQ175 mice and wild type (WT) littermates at the indicated ages. The scale bars are shown on the right and warmer colors represent higher CBVa values. (**e)** Longitudinal CBVa changes in the motor cortex of zQ175 HD mice and WT littermates. Longitudinal data are Mean ± SEM, n = 10. \**p* < 0.05 *versus* the values of the WT group at matched ages by Student’s *t*-test.

We then examined the trajectories of CBVa changes longitudinally in the zQ175 model. CBVa values from multiple ROIs were included as the dependent variables, genotype was included as the independent variable, and genotype-by-time was included as the interaction variable. Significant genotype-by-time interaction was observed between zQ175 mice and controls. We observed that control mice could maintain a stable level of CBVa in basal ganglion regions over the course of 6 months (Figs. 1c, e). In contrast, zQ175 mice exhibited progressively declining CBVa in the striatum [n=10, F(age × genotype)_2,28_ = 6.736,*p*□<□0.01] and motor cortex [F(age × genotype)_2,42_ = 7.046,*p*□<□0.01] over the course of disease progression. Although striatal volumes in this model have been reported before ^38,39^, our measurements were made in the same cohort in which CBVa was measured. This provides more detailed information on possible age-dependent structural changes and more importantly, helps identify the potential influence, of structural changes on CBVa alterations. Our data indicate that zQ175 mice experienced striatal atrophy onset at 6 months (Extended data Fig. 1a, *p* = 0.012) which progressively worsened at 9 months *(p* = 0.001). These HD mice also displayed motor deficits on the tapered beam (Extended data Fig. 1b, *p* = 0.046) and 5 mm balance beam (Extended data Fig. 1c, *p* = 0.018), and decreased locomotor activity in the open field apparatus (Extended data Fig. 1d, *p* = 0.043) at 9 months of age. zQ175 mice also had slower body weight gain than control mice (Extended data Fig. 1e, F(age × genotype)_6,102_ = 8.311,*p*□<□0.01). Taken together, the longitudinal data support that altered CBVa occurs at the premanifest stage, which progressively declines with the onset and progression of manifest HD in the zQ175 model.

### Morphological analysis of the cerebral vasculature in zQ175 mice

Given that mHTT may affect the physiology of cells comprising the cerebral vasculature, we analyzed the morphology and structures of arterioles and cerebral vasculature in zQ175 mice at the premanifest age (3 month) and manifest age (9 month) when significant changes in CBVa were detected between zQ175 mice and controls. Acta2 labels arterioles whereas collagen IV labels all blood vessels including arterioles, capillaries and venules ^41^. At the premanifest stage, there were no significant differences in the vessel density and mean diameter of Acta2^+^ arterioles and collagen IV^+^ blood vessels in the striatum (Fig. 2 a-e) and motor cortex (Extended data Fig. 2 a-e). To compare variability in the diameter of collagen IV^+^ vessels, we used MATLAB software tools to generate skeletonized images ^42,43^ and calculated vascular diameters by executing the Euclidean distance transform ^43^. We located vessel branch points using the skeletonized images and defined a vessel segment as the fragment between two branch points ^44^. Further analysis of blood vessel morphology excluded capillaries (< 2 μm) and focused on smaller vessel diameter ranges at 2-10 μm and 10-20 μm. We did not detect differences in the numbers of vessel segments in the striatum (Fig. 2f-g) and motor cortex (Extended data Fig. 1 f-g) between zQ175 and age-matched controls. These data indicate that elevated CBVa in the premanifest zQ175 mice is not due to permanent morphological changes in cerebral blood vessel densities or diameters, and that it may reflect a transient neurovascular functional response to altered brain energy demand, such as enhanced blood vessel dilation.

**Figure 2.**
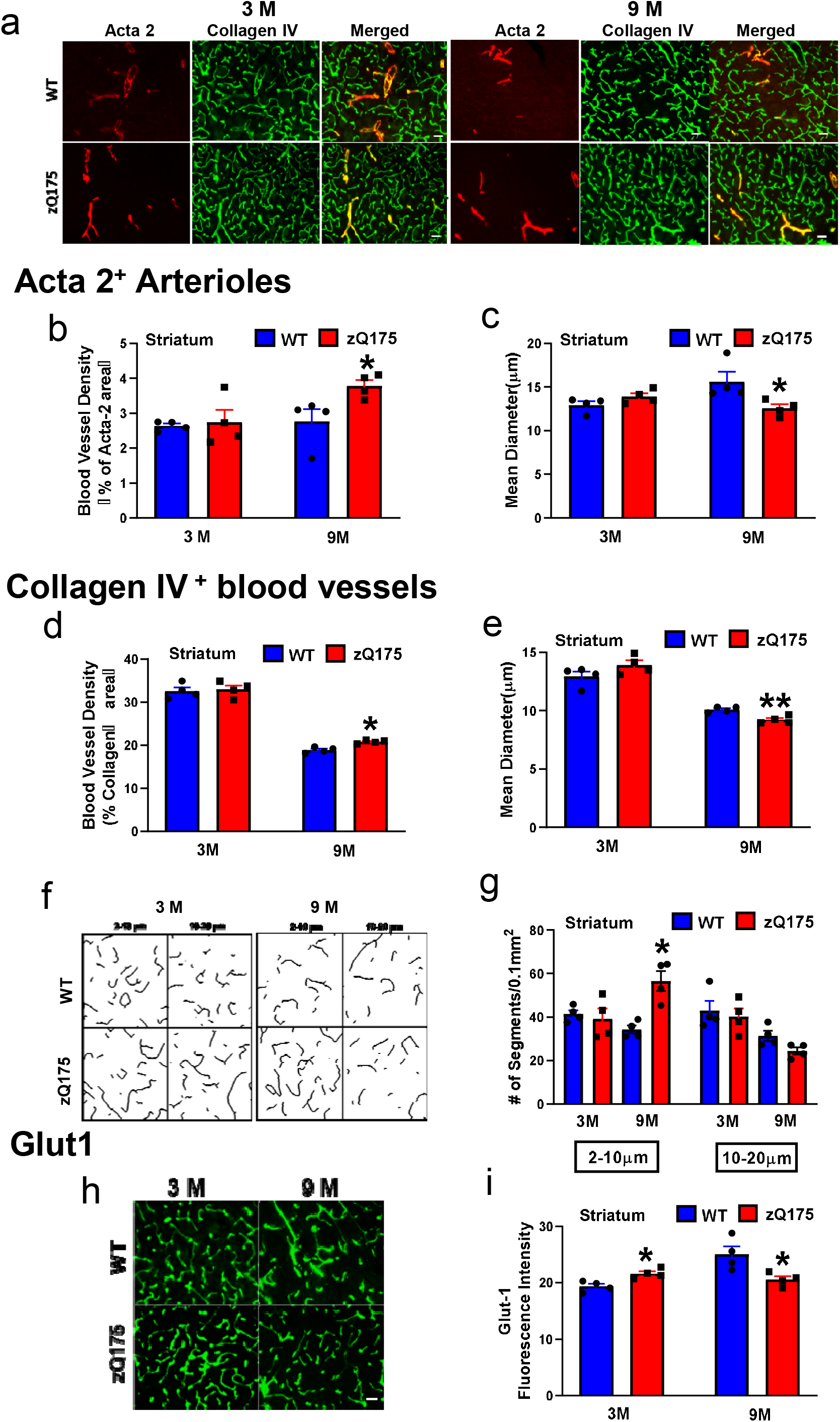
Morphological analysis of cerebral vasculature in the striatum of zQ175 mice. **(a)** Representative images of immunostaining signals of Acta2 (Red, arterioles) and collagen IV (Green, all blood vessels) in the mouse striatum from indicated genotypes and ages. 3 months (3M) represents the premanifest stage in zQ175 mice; 9 months (9M) represents the manifest age of zQ175 mice. Scale bar = 20 μm. (**b-c**) Quantitative analysis of Acta 2^+^ arteriolar density (b) and mean diameter (c) in the striatum of zQ175 mice and wild type (WT) controls at 3 and 9 months of age. (**d-e**) Quantitative analysis of collagen IV^+^ blood vessel density (d) and mean diameter (e) at indicated ages. (**f**) The representative skeletonized image of collagen IV^+^ blood vessels at the indicated diameters in the zQ175 and WT mice at 3 and 9 months of age. (**g**) Quantitative analysis of the number of segments per 2+ 0.1 mm^2^ in the collagen IV blood vessels with indicated diameters. (**h-i**) Representative images of Glut1 immunostaining in the mouse striatum from indicated genotypes and ages (h) and quantitative data of Glut1 immunofluorescent intensity (i). Scale bar = 20 μm. All data are Mean ± SEM, n = 4. \**p* < 0.05 *versus* the values of WT group at the corresponding ages by Student’s *t*-tests.

We next analyzed the morphology of arterioles and cerebral vasculature in manifest zQ175 mice (9 months old) when CBVa significantly declined. At this age, zQ175 mice exhibited increased density (Fig. 2b, *p* = 0.046) and decreased diameters (Fig. 2c, *p* = 0.043) in Acta2+ arterioles in the striatum. Though there still was an increasing trend, no significant differences in arteriolar density in the motor cortex of zQ175 mice were observed (Extended data Fig. 2b, *p* = 0.27). Notably, significantly reduced arteriolar diameter was observed in the motor cortex of zQ175 mice (Extended data Fig. 2 c, *p* = 0.004). Moreover, we observed significantly increased density (Fig. 2d, *p* **=** 0.009) and reduced vessel diameter (Fig. 2e, *p* = 0.005) in all collagen IV^+^ striatum vasculatures of 9-month-old zQ175 mice. The effects of increased density and reduced vessel diameter were smaller in collagen IV^+^ vessels compared to Acta2^+^ arterioles in the striatum of 9-month-old zQ175 mice. Further analysis by separating collagen IV^+^ blood vessels into different groups based on diameter as above for 3-month-old mice showed that the number of segments in smaller blood vessels (diameter at 2-10 μm) was significantly higher (Fig. 2f-g, *p* = 0.004) in the striatum of zQ175 mice, while no significant changes in the numbers of larger blood vessel segments (diameter at 10-20 μm) (Fig. 2f-g) were observed. Similar changes in the morphology of collagen IV^+^ blood vessels were also detected in the motor cortex of zQ175 mice (Extended data Fig. 2f-g). These results suggest that lower CBVa in manifest HD mice may be at least partially due to these permanent morphology changes of blood vessels, particularly reduced arteriolar diameter. These structural changes in the collagen IV^+^ cerebral blood vessels of manifest zQ175 mice were consistent with reported vascular alterations in other HD mouse models ^36,45^. Because the cerebral vascular system provides essential support for effective brain functioning ^46^, morphological abnormalities in the cerebral vasculature likely further compromise neuronal integrity and function, thereby contributing to HD pathology and disease progression.

CBVa is a hemodynamic measure that may be an indirect indicator of neuronal metabolism and function ^47^, as altered neuronal activity and/or glucose utilization can result in differences in CBVa readouts. It is worth noting that blood vessel diameters measured from histology may not reflect the blood vessel diameters measured during *in vivo* MRI scans when the animals are alive. Because we did not find histological changes in the density and diameter of arterioles and cerebral blood vessels in premanifest zQ175 mice (3 month), we hypothesize that elevated CBVa measured with iVASO MRI at the premanifest HD stage may represent a compensatory vascular dilation in response to impaired neuronal metabolism. If this is the case, glucose transporters may also be upregulated to meet the neuronal energy demands of the premanifest HD brain. We thus conducted immunofluorescent staining to examine the major glucose transporter Glut1 in endothelial cells. Significantly increased Glut1 immunofluorescent intensity was observed in the striatum (Fig. 2h-i, *p* = 0.033) and motor cortex (Extended data Fig. 2 h-i, *p* = 0.020) of premanifest zQ175 mice (3M), suggesting upregulated glucose transport in the HD brain. We next asked whether such upregulation of the glucose transporter will persist to the manifest stage. In contrast, manifest zQ175 HD mice (9M) exhibited significantly decreased Glut1 immunofluorescent intensity in the striatum (Fig. 2 h-i, *p* = 0.049) and motor cortex (Extended data Fig. 2 h-i, *p* = 0.036). Taken altogether, our data suggest that the HD brain initiates a compensatory cerebral vascular response to altered neuronal activity and/or energy metabolism in the premanifest stage, while impaired vasculature structure leads to lowered CBVa and possibly compromised compensatory regulation - such as reduced glucose transporter levels-thus exacerbating HD pathology and manifestation.

### CRISPR/Cas9-mediated *HTT* silencing in the striatal neurons restores altered CBVa in premanifest zQ175 mice

We next tested whether suppression of mHTT in neurons could normalize altered CBVa in the premanifest HD brain. Lowering mHTT by introducing CRISPR/Cas9-mediated HTT silencing to the striatal neurons has been reported to improve motor function in HD mice with rare off-target effects ^31^. As demonstrated in our longitudinal characterization data, heterozygous zQ175 mice do not show motor phenotype and striatal atrophy at 3 months of age (Extended data Fig. 1). Therefore, *HTT* silencing agents were administered into the striatum of 2-month-old mice. Outcome measures, including CBVa, striatal volume, and motor function, were assessed one month later.

To lower mHTT efficiently, we used combinations of two guiding RNAs targeting human *HTT* exon-1 regions (Fig. 3a, T1 and T3, designed by Xiaojiang Li’s group at Emory then). As T1 gRNA recognizes mouse *Htt* exon-1 and T3 gRNA exclusively recognizes human *HTT* exon-1, combinations of T1 and T3 could more efficiently lower mHTT (human exon-1) levels than those of wild type HTT (mouse exon-1). Two gRNAs are expressed under the U6 promoter in an AAV vector that also expresses red fluorescent protein (RFP) (AAV-HTT gRNAs) for the purpose of tracking expression.

**Figure 3.**
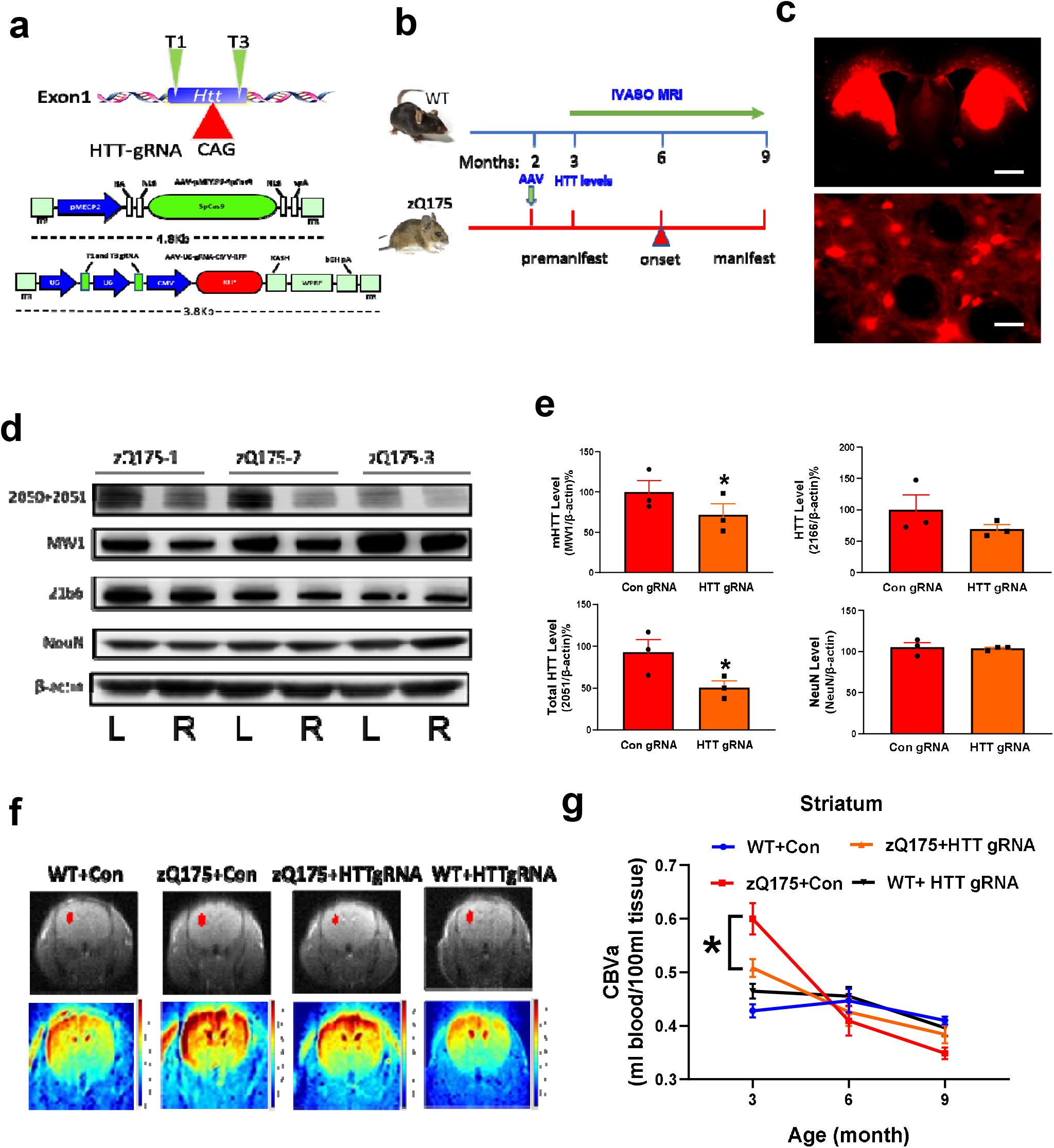
CRISPR/Cas9-mediated *HTT* silencing in the striatal neurons restores altered CBVa in the striatum of zQ175 mice. (**a**) Schematics of the designed HTT-gRNA (T1 and T3) and AAV vectors. ITR, inverted terminal repeat; HA, human influenza hemagglutinin; NLS, nuclear localization sequence; KASH, Klarsicht, ANC-1, Syne Homology; WPRE, woodchuck hepatitis virus post-transcriptional regulatory element; RFP, red fluorescent protein. (**b**) Timeline for AAV injections, HTT level measures and longitudinal CBVa assessment. (**c**) RFP fluorescence showing the transduction of AAV-HTT-gRNAs in the striatum. Scale bar = 900 μm (top image) and 40 μm (bottom image). (**c**) The images of Western blotting with indicated antibodies to HTT, NeuN, and β-actin. (**e**) Quantification results of mHTT (MW1), total HTT (combining upper and lower bands in the MCA2050 + MCA2051 blot), wtHTT (lower and major band in the blot with MAB 2166 antibody), and NeuN in the striatum injected with AAV-HTT gRNA + Cas9 (HTT gRNA, right striatum-R) or AAV-control gRNA + Cas9 (Con gRNA, left striatum-L). n=3, \**p* < 0.05 *versus* the values of con gRNA group by Student’s *t*-tests. (**f**) Representative CBVa maps in mouse brains from indicated genotypes and treatment groups. Top row shows the raw images, and the red ROIs indicate the quantified brain region. Bottom row shows the representative CBVa maps in the mice at the indicated genotypes and treatment at 3 months of age. The scale bars are shown on the right and warmer color represents higher CBVa values. (**g**) Quantification of longitudinal CBVa changes in the striatum of zQ175 HD mice and WT littermates with indicated treatment at indicated ages. Mean ± SEM, n = 6-7. * *p* < 0.05, comparison between zQ175 mice injected with HTT gRNAs (+ spCas9) *versus* zQ175 mice injected with control RNA (+ spCas9) by Two-way *ANOVA* with Bonferroni *post hoc* analysis.

The knock-down efficiency of these combined HTT gRNAs has been demonstrated previously ^31^. spCas9 is expressed in a separate AAV vector under the Mecp2 promoter (AAV-Mecp2-spCas9) ^31^ in order to drive the construct to express in neurons only.

To confirm the efficacy of CRISPR/cas9-mediated approach to lower HTT in zQ175 mice. We designed experiments as indicated (Fig. 3b). AAV_9_-HTT gRNAs and AAV-Mecp2-spCas9 were injected into one side of the striatum (right side, R) and the contralateral striatum (left side, L) was injected with AAV9-control gRNA and AAV-Mecp2-spCas9. The two viruses were mixed at a ratio of 1:3 for stereotaxic injection. RFP fluorescence (gRNA construct contains) in the striatum indicated successful injection and virus transduction (Fig. 3c). Western blotting with different HTT antibodies indicated that levels of mutant HTT (upper band in the 2050+2051 blot, MW1 blot) and wild type HTT (strong band in the 2166 blot and lower band in the 2050+2051 blot) were significantly reduced by HTT gRNA + spCas9 injection (Fig. 3d-e) 4 weeks after injection. NeuN (a pan neuronal marker) levels (Fig. 3 d-e) were examined to verify that reduced HTT levels were not due to non-specific pan neuronal damage in the AAV-injected striatum.

We then evaluated the effects of lowering HTT in striatal neurons at the premanifest stage on altered CBVa in zQ175 mice. HTT gRNAs (or control) + spCas9 were injected into the bilateral striatum of 2-month-old mice and outcomes were assessed at 3 months of age. Remarkably, zQ175 mice injected with HTT gRNA (and spCas9) in the striatum had rescued elevated CBVa in the striatum at 3 months (Fig 3 f,*p* < 0.05). Mice injected with HTT gRNA + spCas9 in the striatum exhibited a CBVa curve close to normal mice [Fig. 3g, WT+ Con *vs* zQ175 + Con, F(age × Genotype)_2,24_ = 19.51,*p*□< 0.01; zQ175 + Con *vs* zQ175 + HTT gRNA, F(age × Treatment)_2,24_ = 4.7, \**p*□ < 0.05]. Lowering HTT in striatal neurons did not significantly affect the altered CBVa in the motor cortex (Extended data Figs. 3 a-b). These results suggest that the elevated CBVa in premanifest HD is most likely due to neuronal changes in either activity or metabolism within the brain regions we measured. Our data confirmed that this non-allele-specific HTT lowering did not affect body weight or locomotor activity in control mice as well as zQ175 mice (Extended data Figs 3 c-d). Because the iVASO MRI technique has been developed and successfully applied to the human brain ^48^, the present study provides a proof of principle to consider CBVa measured by iVASO as a biomarker to evaluate therapeutic efficacy in premanifest HD clinical trials.

### *HTT* silencing at the premanifest stage delays the onset and slows the progression of HD-like phenotype and pathology in zQ175 mice

Given that the onset and severity of clinical manifestations in HD does not only depend on neuronal loss but also on neuronal dysfunction and circuitry reorganization, these processes may occur at an early stage of the disease, possibly before neurodegeneration ^49^. We asked whether lowering HTT at the premanifest stage could delay or even prevent the onset of HD. HTT gRNAs (or control) + spCas 9 were administered to zQ175 mice and control mice at 2 months of age. Motor function and striatal volume were assessed longitudinally from 3 months to 9 months (Fig. 4a). Similar to untreated zQ175 mice, control gRNAs + spCas9 injected zQ175 mice displayed significant striatal atrophy (Fig. 4b, *p* < 0.01) and motor impairment (Fig. 4c-d, *p* < 0.05) at 6 months. In contrast, zQ175 mice treated with HTT gRNAs + spCas9 had no significant striatal atrophy (Fig. 4b) or motor deficits (Fig. 4c-d) at 6 months, indicating that HTT lowering at the premanifest stage delays the onset of motor phenotype and striatal atrophy in zQ175 HD mice.

**Figure 4.**
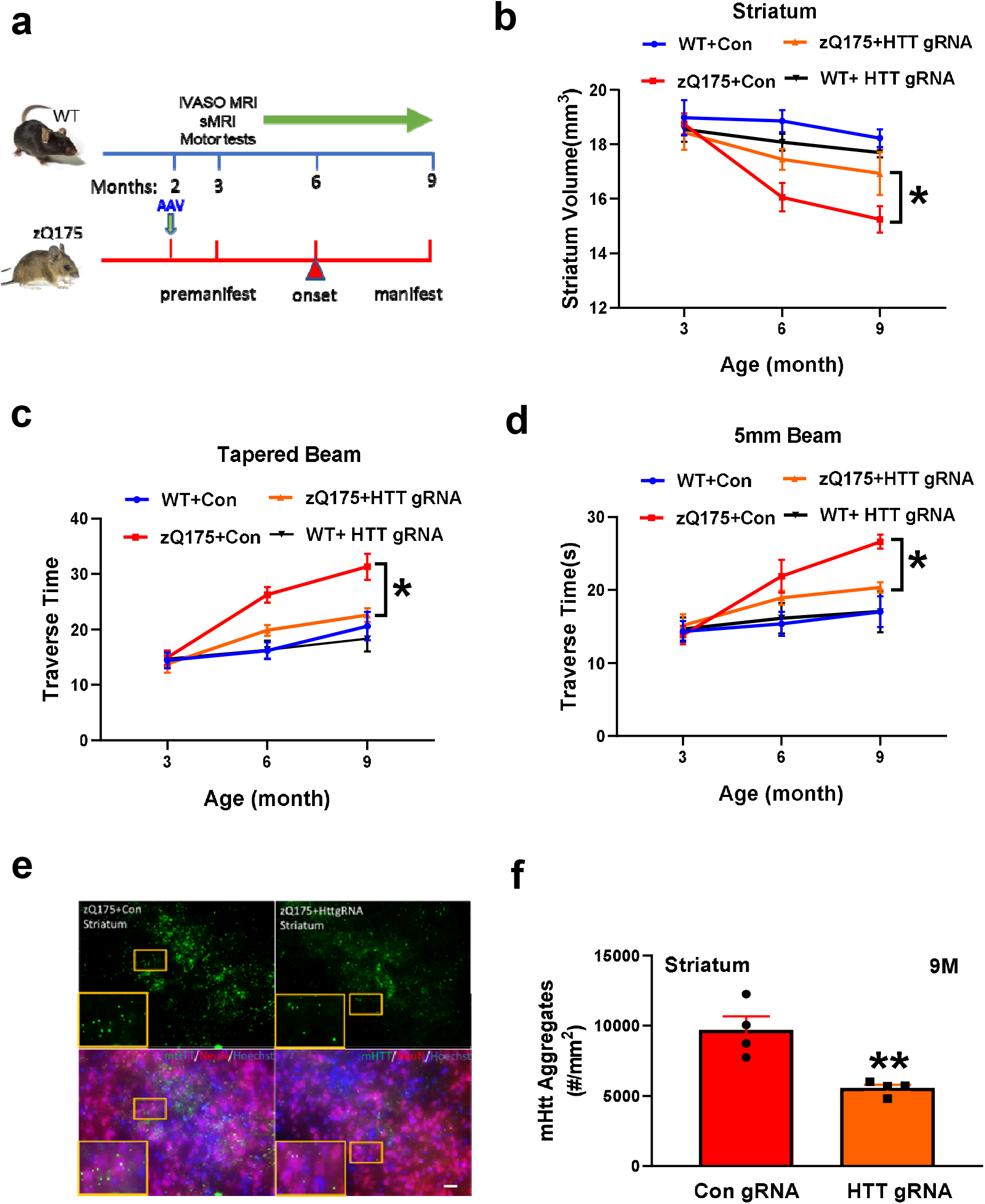
*HTT* silencing at the premanifest stage delays the onset and slows the progression of HD-like phenotype and pathology in zQ175 mice. (**a**) Timeline for AAV injections and outcome measures. **(b)** Longitudinal striatal volume data were quantified from structural MRI scans from indicated groups at indicated ages. (**c**) Mice were tested on a tapered beam and time crossing the beam (Traverse time) was recorded from different groups at indicated ages. (**d**) Mice were tested on a 5 mm beam and time crossing the beam (Traverse time) was recorded from different groups at indicated ages. All data in (b) through (d) are Mean ± SEM, n = 6-9. \**p* < 0.05, comparison between zQ175 mice injected with HTTg RNAs (+ spCas9) *versus* zQ175 mice injected with control RNA (+ spCas9) by Two-way *ANOVA* with Bonferroni *post hoc* analysis. (**e**) Mutant HTT (mHTT) aggregates were labeled by immunostaining with EM48 antibody in the striatum of zQ175 mice at 9 months of age. EM48-positive mHTT aggregates (green), pan neuronal marker NeuN (red), and nucleus (blue, DAPI) were indicated. Scale bar = 20 μm. (**f**) Numbers of mHTT aggregates (per mm^2^) were quantified in the striatum. Mean ± SEM, n = 4, \*\**p* < 0.01 *versus* the values of con gRNA group by standard Student’s *t*-tests.

We then assessed whether lowering HTT at the premanifest stage slowed the progression of manifest HD by comparing longitudinal data. Remarkably, *HTT* silencing in the striatal neurons significantly slowed the progression of striatal atrophy [Fig. 4b, WT+ Con *vs* zQ175 + Con, F(age × Genotype)_2,24_ = 6.926,*p*□ < 0.01; zQ175 + Con *vs* zQ175 + HTT gRNA, F(age × Treatment)2,22 = 3.584, #*p*□< 0.05] and ameliorated motor deficits on both tapered beam [Fig. 4c, WT + Con *vs* zQ175 + Con, F(age × Genotype)_2,28_ = 5.562, *p*□ < 0.01; zQ175 + Con *vs* zQ175 + HTT gRNA, F(age × Treatment)_2,40_ = 2.1, \**p*□< 0.05] and 5 mm beam tests [Fig. 4d, WT + Con *vs* zQ175 + Con, F(age × Genotype)2,27 = 4.635,*p*□ < 0.05; zQ175 + Con *vs* zQ175 + HTT gRNA, F(age × Treatment)_2,35_ = 3.382, \**p*□< 0.05] in zQ175 mice. CBVa levels in the motor cortex, body weight, and locomotor activity were not significantly modulated by lowering HTT in the striatum (Extended Fig. 3 a-d).

A common histopathological hallmark of HD brains is the presence of pathognomonic mHTT aggregates, though its role in disease pathology is still unclear and controversial. We determined whether HTT silencing affected mHTT aggregation, as mHTT aggregates may form due to mutant HTT that increases the cellular concentrations of disease-specific proteins or renders them aggregation-prone. Thus, HTT silencing may reduce the mHTT aggregates if mHTT is efficiently lowered. After the last longitudinal assessment at 9 months, zQ175 mice were sacrificed and mHTT aggregates were labeled by immunostaining with the EM48 antibody (Fig. 4e). We found that HTT gRNA + Cas9-injected zQ175 mice had significantly less mHTT aggregates in the striatum compared with mice injected with control gRNAs (Fig. 4f, ** *p* < 0.01), suggesting that this HTT silencing strategy efficiently lowered mHTT levels.

## Discussion

A challenge in developing treatments for HD is the identification of demonstrable outcome measures for evaluating intervention efficacy in delaying onset or improving underlying pathological processes prior to measurable clinical symptoms. Our study has shown for the first time that a sensitive physiological MRI measure responds to an HTT-lowering treatment in the premanifest period of HD, and demonstrates the feasibility of detecting arteriolar perfusion changes in the premanifest HD brain with a robust MRI technique that is suitable for longitudinal evaluations of therapeutic efficacy. Moreover, we provide the first evidence that introducing HTT-lowering treatment before the occurrence of motor symptoms and striatal atrophy delays onset and slows progression of HD-like phenotype manifestation in a full-length HD mouse model. The present results also suggest that elevated CBVa in premanifest HD is secondary to the effects of mHTT on neural activity/metabolism, and that a reduced rate of nutrient delivery due to reduced CBVa and decreased glucose transporter Glut1 across a compromised neurovascular network in manifest HD might become the primary event that eventually triggers neuronal dysfunction and degeneration.

Rodent HD models play an important role in elucidating phenotypic progression and in investigating potential underlying mechanisms of clinical symptoms ^50^. The validation of potential biomarkers in animal models has aided in the preclinical development of new therapeutic modalities. However, neuroimaging, especially functional and physiological brain imaging methods, has only recently begun to be applied to HD mouse models. In the present study, we utilized iVASO MRI, developed initially for the human brain, and observed similar alterations in premanifest mouse and prodromal human HD brains. Our findings demonstrate that significant changes in CBVa occur before striatal atrophy and motor symptoms, further supporting that altered cerebrovascular function is an early event in HD.

The fact that CBVa changes emerge over time in the striatum and motor cortex supports that dysfunction in the basal ganglion circuitry is associated with disease phenotype, though we do not know the underlying mechanisms that link CBVa and neuronal dysfunction (or compensation for dysfunctional neuronal metabolism) in premanifest HD. However, these changes indicate that there is a significant therapeutic window to test interventions in full-length HD mouse models, such as the heterozygous zQ175 model^39^, which is not present in faster-progressing models. While no animal model replicates all the features of HD, the heterozygous zQ175 HD model offers an alternative system to study functional changes in the premanifest HD brain.

The use of iVASO MRI for CBVa assessment may provide an improved means for the bidirectional translation of biomarkers between animal models and human HD. Reverse translation of biomarkers to rodents also provides an important tool for evaluating the therapeutic effects of candidate treatment ^51^, which is particularly relevant as several clinical trials on lowering HTT using different strategies, including ASO and RNA interference, are ongoing ^4^. It is notable that most HD mouse models including the zQ175 model are not characterized by overt neuronal death, which argues against a direct involvement of neuronal loss. Such interventions may target neuronal dysfunction and consequently modify disease progression. In fact, the severity of clinical manifestations in HD likely does not depend solely on neuronal loss but also on neuronal dysfunction and circuitry reorganization ^49^. Our data supports the existence of cerebrovascular abnormality in HD that may be independent from the neuronal loss characteristic of the disease. Overall, this study not only reveals a promising imaging biomarker for premanifest HD, but also uncovers potential novel therapeutic windows. Future studies will need to address the predictive validity of using CBVa in HD clinical trials and giving HTT-lowering treatment in premanifest HD.

Microvascular abnormalities in different segments of the microvasculature (arteriolar and venous blood vessels) have been identified in neurodegenerative diseases ^8,20^. The supply of adequate oxygen and energy substrates for local metabolic demands is controlled by blood vessels and the status of the microvasculature is closely associated with brain functions and energy metabolism. Different types of blood vessels have distinct functions and physiology and may be differentially affected by HD pathology. Arterioles are the most actively regulated blood vessels, and thus may be more sensitive to metabolic disturbances in the brain^28–30^. The iVASO MRI approach can measure CBV in small pial arteries and arterioles (CBVa) ^28,48^, the primary regulator of local tissue perfusion. The iVASO method has been used to measure neurovascular abnormalities in human brains of several disease conditions, such as brain tumors ^52^, HD ^15^, Parkinson’s disease ^53^, Alzheimer’s disease dementia ^53,54^, and schizophrenia ^55^. In the current study, we demonstrated that CBVa was significantly increased at premanifest stage in heterozygous zQ175 mice, while structural MRI measures showed significant striatal atrophy later, indicating that iVASO CBVa measures may be a sensitive biomarker for assessing therapeutic response in the premanifest HD clinical trials.

Current ongoing clinical trials of HTT-lowering approaches target all cells in the brain. Our study for the first time demonstrated that *HTT* silencing in striatal neurons can restore CBVa in the basal ganglion circuitry of premanifest HD brains, consequently delaying the onset of striatal atrophy and motor deficits, and slowing pathological progression of HD. Our results implicate that elevated CBVa in premanifest HD is likely a compensatory response to altered energy demands in striatal neurons, as reducing mHTT levels in those neurons rescued the elevated CBVa in the striatum of premanifest HD mice. This early compensatory response could be impaired over the course of disease progression due to permanent morphological changes in the cerebral vasculature during the manifest HD period. Thus, an inability to increase blood supply in response to increased energy demands may accelerate neuronal dysfunction and degeneration in the long term.

An underlying assumption in our study was that CBVa precisely reflected specific pathophysiological processes. However, we are aware that iVASO MRI techniques still offer a limited characterization of vascular properties. Consequently, it is important to exert caution about the observed CBVa changes with disease progression. Although abnormal cerebrovascular trajectories may reflect a tentative ordering in which pathophysiological events occur, our results should be interpreted within the scope of biomarker sensitivity to disease progression instead of causal pathologic interactions. Another potential limitation of our study is that all evaluations were performed within a linear regression framework, which could mean that obtained results primarily reflect linear tendencies in the analyzed outcome measures. These traditional limitations suggest the need to study disease progression not only in terms of alterations of specific biomarkers, but also through analysis of the multifactorial causal pathological interactions that take place on different spatiotemporal scales.

There is increasing evidence that a number of neurodegenerative disorders are associated with alterations in the cerebral vasculature ^46^. A multifactorial data-driven analysis for all biomarkers and a tentative temporal ordering of disease progression indicate that cerebrovascular dysregulation is an early pathological event during Alzheimer’s disease (AD) development ^56^. Our present findings implicate a similar pathology in HD. The highly regulated neurovascular coupling system ensures that regional blood supply is increased to meet energy needs and remove metabolic waste. This system often is impaired by disease pathology. Our data suggest that elevated CBVa may be one of the initial steps contributing to observed vascular modifications in premanifest HD and restoration of CBVa during this particular period by the HTT-lowering approach or other disease modifying treatment could be an indication of improved neuronal function/metabolism. Additionally, we demonstrate that early longitudinal measures of CBVa changes may be predictive of later therapeutic efficacy, and that lowering HTT before clinical symptoms manifest has long-lasting beneficial effects in HD. Further validation of these findings in human clinical trials will facilitate the development of efficient therapeutic interventions for premanifest HD, with a goal to delay or even conceivably prevent onset of manifest HD.

## Materials and Methods

### Animals

Heterozygous zQ175 mice (both males and females) and wild type littermates were used in the study, and the zQ175 breeders were purchased from the Jackson Lab (Bar Harbor, ME). Genotyping and CAG repeat count were determined by PCR of tail snips at Laragen Inc. (Culver City, CA, USA). The CAG repeat length was 220 ± 3 in the zQ175 mice used in the study. All mice were housed at 3-5 mice per cage under specific pathogen-free conditions with a reversed 12-h light/dark cycle maintained at 23°C and provided with food and water ad libitum. All behavioral tests and longitudinal measures were done in the mouse dark phase (active). The study was carried out in strict accordance with the recommendations in the Guide for the Care and Use of Laboratory Animals of the National Institutes of Health and approved by Institutional Animal Care and Use Committee at the Johns Hopkins University. The protocol was approved by the Committee on the Ethics of Animal Care and Use Committee (Permit Number: MO18M192). All MRI procedures were performed under isoflurane anesthesia, and all efforts were made to minimize suffering.

### MRI acquisition

*In-vivo* studies were performed on a horizontal bore 11.7 T Bruker Biospec system (Bruker, Ettlingen, Germany) equipped with a physiological monitoring system. A 72-mm quadrature volume resonator was used as a transmitter, and the receiver was a four-element (2 × 2) phased-array coil. Anatomical images covering the entire brain were acquired using a multi-slice Rapid Acquisition using Relaxation Enhancement (RARE) sequence: resolution=0.10× 0.10 × 0.50 mm^3^, echo time (TE)/repetition time (TR)=30/3000 ms, and RARE factor = 8. A multi-slice iVASO sequence optimized for typical blood flow and vascular transit times in mice and relaxation times at 11.7T was performed for CBVa mapping: TR/inversion time (TI) = 1924/800, 1636/700, 1365/600, 1109/500, 867/400, 750/350, 636/300, 525/250, 415/200 ms, TE = 4.6 ms, resolution=0.15×0.15×0.80 mm^3^, and RARE factor=16. All mice were anesthetized with an induction of 2% isoflurane in medical air, and then 1.0-1.5% isoflurane during the MRI scan. The mouse head position was fixed with a bite bar and two ear pins. During MRI scans, the animal was placed on a water-heated animal bed equipped with respiratory controls. The animal’s respiration rate was constantly monitored using an animal monitoring system (SAII, Stony Brook, NY, USA). Mice were ventilated to maintain stable physiologic conditions (respiratory rate at 60-80 breaths/min).

### iVASO MRI image analysis

Statistical parametric mapping (SPM) (Version 8, Wellcome Trust Centre for Neuroimaging, London, United Kingdom; http://www.fil.ion.ucl.ac.uk/spm/) and in-house programs coded in Matlab (MathWorks, Natick, MA, USA) were used. Motion correction was performed for all iVASO images, and the difference in iVASO signal was calculated using the surround subtraction method^57^. Cortical thicknesses obtained from structural MRI scan were used to correct partial-volume effects on the iVASO signal. Whole-brain CBVa maps were calculated using the iVASO theory ^48^. Two regions-of-interest (ROIs) were manually delineated in each scan: 1) motor cortex, and 2) striatum. Average CBVa values were obtained in each ROI.

### Structural MRI image analysis

Images were first rigidly aligned to a template image by using automated image registration software (http://bishopw.loni.ucla.edu/AIR5/, AIR). The template image was selected from one of the images acquired from age-matched littermate control mice (mouse had the medium brain volume among the control group), which had been manually adjusted to the orientation defined by the Paxinos atlas with an isotropic resolution of 0.1 × 0.1 × 0.1 mm^3^. After rigid alignment, images had the same position and orientation as the template image, and image resolution was also adjusted to an isotropic resolution of 0.1 × 0.1 × 0.1 mm^3^. Signals from non-brain tissues were removed manually (skull-stripping). Skullstripped, rigidly aligned images were analyzed by the Landmarker software (www.mristudio.org). Intensity values of the gray matter, white matter, and cerebral spinal fluid were normalized to values in the template images by using a piece-wise linear function. This procedure ensured that subject image and template image have histograms of similar intensity. The intensity-normalized images were submitted by Landmarker software to a Linux cluster, which runs Large Deformation Diffeomorphic Metric Mapping (LDDMM). The transformations were then used for quantitative measurement of changes in local tissue volume among different mouse brains, by computing the Jacobian values of the transformations generated by LDDMM.

### Behavioral tests

<u>5mm balance beam testing</u> was conducted on an 80-cm long and 5-mm wide square-shaped balance beam that was mounted on supports of 50-cm in height. A bright light illuminated the start platform, and a darkened enclosed 1728 cm^3^ escape box (12 × 12 × 12 cm^3^) was situated at the end of the beam. Disposable pads placed under the beam provided cushioning if an animal fell off. Mice were trained to walk across the beam twice at least 1 h prior to testing. If a mouse stopped during training, the tail was gently pressed to encourage movement. After the training trial, mice were left undisturbed for at least an hour before testing. The time for each mouse to traverse the balance beam was recorded with a 125-sec maximum cut-off, and falls were scored as 125 sec.

<u>Tapered beam testing</u> was conducted on a beam that is 1 m in length tapering from 3.5 cm to 0.5 cm with underhanging ledges 1.0 cm in width on either side. The beam was placed at a 30° angle of incline, with the narrowest end at the highest point. Animals were pre-trained for 3 trials in order to habituate them to the task, and then tested at indicated ages. After the training trial, mice were left undisturbed for at least an hour before testing. The time for each mouse to traverse the tapered beam was recorded with a 125-sec maximum cut-off, and falls were scored as 125 sec.

<u>Open field locomotor activity</u> was performed during the dark phase of the diurnal cycle under red light conditions. The open field tests were performed at the ages of 3 months, 6 months, 9 months and 12 months old mice. The mice were housed in the experimental room. Locomotor activity was measured by an automated Open Field Activity System and the data were analyzed by Activity Monitor software (Columbus Instrument Inc., OH). The activity chambers (27.3 × 27.3 × 20.3 cm^3^) were equipped with infrared beams. Mice were placed in the center of the chamber and their behaviors were recorded for 60 min with 5-min bins for total activity, peripheral and central activity, as well as the rear frequency.

### AAV reagents and stereotaxic injection

CRISPR/Cas9-related viral vectors (PX551, PX552) were kindly provided by Dr. Xiaojiang Li’s lab (Emory University, then). The original viral vector was obtained from Addgene (plasmids #60957 and 60958). AAV_9_-HTT-gRNAs were generated by inserting gRNAs into PX552 via SapI restriction sites. gRNA sequences are as follows: T1: GGCCTTCATCAGCTTTTCCAggg, T3: GGCTGAGGAAGCTGAGGAGGcgg, and control gRNA: ACCGGAAGAGCGACCTCTTCT (PAM sequence is shown in lowercase). AAV_9_-Mecp2-Cas9 vector was generated in PX551 with CMV promoter (658 bp) using XbaI and AgeI restriction sites. These viral vectors were sent to the Viral Vector Core at Emory University for packaging and purification of viruses. The genomic titer of viruses was determined by PCR method.

The mice were anesthetized with 1.5% isoflurane inhalation and stabilized in a stereotaxic instrument (David Kopf Instruments). All surgical procedures were performed in a designated procedure room and in accordance with the Guidelines for the Animal Care and Use Committee and biosafety procedures approved by the Johns Hopkins University School of Medicine. All mice were injected unilaterally or bilaterally into the striatum using the stereotaxic coordinates: 0.62 mm rostral to bregma, ±1.75 mm lateral to midline and 3.5 mm ventral to the skull surface. zQ175 mice and wild type littermates were injected with AAV_9_ expressing gRNA and spCas9 which were mixed at a ratio of 1:3, and 2 μl of the mixed viruses (1.0-1.3 × 10^13^ particles/μl) were injected into each side of the mouse striatum using a Hamilton syringe and a syringe infusion pump (World Precision Instruments, Inc., Sarasota, Florida, USA). Small holes were drilled in the skull, and a 30-gauge Hamilton micro syringe was used to deliver the virus at a perfusion speed of 0.2 μl/min.

### Immunohistochemistry

Mice were anesthetized and perfused transcardially with phosphate-buffered saline (PBS) followed by 4% paraformaldehyde. Brains were post-fixed overnight followed by immersion in 30% sucrose for 24 h. Coronal brain sections (40 μm) were cut on a cryostat. Sections were stained with primary antibodies, including EM48 (MAB5347, 1:200, Merck Millipore, USA), NeuN (26975-I-AP, 1:200, Proteintech, USA), Collagen □ (2150-1470, 1:100, BioRad, USA), Acta-2(C6198, 1:200, Millipore Sigma, USA), and GLUT1 (SPM498, 1:200, Thermo Fisher, USA). Briefly, the sections were washed three times with PBS for 10 min each time, then permeabilized by incubating with 0.3% Triton X-100 for 5 min, followed by incubation with blocking solution containing 5% donkey serum, 3% goat serum and 0.3% Triton X-100 for 1 h. The sections were then incubated with primary antibody at 4°C overnight. After three washings with PBS, the sections were incubated with fluorescence-labeled secondary antibody for 2 h at room temperature, and then washed 3 times with PBS. Sections were mounted onto Superfrost slides (Fisher Scientific, Pittsburgh, PA, USA) dried and then covered with anti-fade mounting solution. Fluorescence images were acquired with Keyence BZ-X700 All-in-One florescence microscope.

### Blood vessel analysis

After immunofluorescence images were acquired, we used Matlab Image Processing ToolboxTM which provides a comprehensive set of image processing and morphological analysis algorithms to perform image contrast adjustment, noise reduction and segmentation. In order to reduce processing time and enhance signal-to-noise ratio, we converted RGB true color image to grayscale. We then used the imaging adjusting tool to increase contrast in low-contrast areas and sharpen differences between black and white which can help to identify the blood vessels and tissues. For the purpose of smoothing out the noise in the background and normalizing the vessel intensities, a 2-D Gaussian filter was applied ^58^. To quantify vessel structure, Bradley’s method ^59^ was used to create a binarized image. This method can compute the threshold for each pixel using an adaptive technique. We then generated a skeletonized image by using a morphological method ^42,43^, and calculated the vascular diameters by executing the Euclidean distance transform ^43^. We could locate the branch points using the skeletonized image. Vessel segment was defined as the fragment between two branch points ^44^. Thereafter, the segment length was decided by the linear distance between start and end points. Total vascular length was calculated by summing the distances of all segments. Blood vessel density was quantified as the percentage of stained area in the whole visual field.

### Western Blotting

Striatal tissue samples were homogenized in a buffer containing 50 mM Tris-HCl, pH 8.0, 150 mM NaCl, 0.1% (w/v) SDS, 1.0% NP-40, 0.5% sodium deoxycholate, and 1% (v/v) protease inhibitors. For SDS PAGE, 30–50 μg of proteins were separated in a 4-20% gradient gel and transferred to a nitrocellulose membrane. The membrane was blotted with the following primary antibodies: HTT (MAB 2166, 1:1000, MilliporeSigma, USA), HTT (MCA2050, 1:1000, BioRad, USA), HTT (MCA2050, 1:1000, BioRad, USA), mHTT (MW1, Anti-poly-Q, 1:1000, Millipore Sigma, USA) and mouse anti-β-actin (Sigma, mouse monoclonal antibody, 1:5000). After incubation with HRP-conjugated secondary antibodies, bound antibodies were visualized by chemiluminescence. The intensity of the Western blot bands was quantified by the ImageJ software.

### Statistics

Data are expressed as the mean □±□ SEM unless otherwise noted. Statistical analysis was performed with SPSS using two-tailed Student’s *t*-test, one-way ANOVA, two-way ANOVA, with Bonferroni *post-hoc* tests as indicated. Graphs were prepared using GraphPad Prism. The *p*-values less than 0.05 were considered as statistically significant. N is reported in the figure legends.

### Data Availability

The authors confirm that all the data supporting the findings of this study are available within the article and its Supplementary material. Raw data will be shared by the corresponding author on request.

## Acknowledgement

This work is supported by R21NS104480 from NINDS (to J.H. & W.D). We thank other financial supports from NINDS R01NS082338 (to W.D). We thank Drs. Xiaojiang Li and Shihua Li (at Emory University then) for providing both HTT gRNA and CRISPR/Cas9 constructs.

## Author contributions

H.L. designed and conducted the experiments, interpreted the data, and contributed to manuscript writing. C. Z. conducted iVASO MRI scans and imaging analysis, behavioral tests and data analysis, and morphological analysis of cerebral vasculature. J.X. designed the iVASO MRI sequence for mouse scanner and directed the iVASO MRI scans and imaging acquisition. J.J. assisted the EM48 immunostaining and imaging analysis. L.C. contributed to imaging analysis, behavioral tests, Western blotting experiments, and manuscript editing. Q.W. contributed to behavioral tests and data analysis. Z.W., P.L. and H.L. assisted animal physiological parameter assessment during iVASO MRI scans. P.C.M.V.Z. contributed to iVASO technique development, data interpretation, and manuscript editing. C.A.R contributed to data interpretation, discussion and manuscript writing. J.H. developed the iVASO fMRI technique and co-designed the project and contributed to manuscript writing. W.D. designed, directed, and coordinated the project and wrote the paper.

## References

1. Ross, C.A., et al. Huntington disease: natural history, biomarkers and prospects for therapeutics. Nat Rev Neurol 10, 204–216 (2014).

2. Bates, G.P., et al. Huntington disease. Nat Rev Dis Primers 1, 15005 (2015).

3. Ross, C.A., et al. Movement Disorder Society Task Force Viewpoint: Huntington’s Disease Diagnostic Categories. Mov Disord Clin Pract 6, 541–546 (2019).

4. Tabrizi, S.J., Ghosh, R. & Leavitt, B.R. Huntingtin Lowering Strategies for Disease Modification in Huntington’s Disease. Neuron 101, 801–819 (2019).

5. Wild, E.J. & Tabrizi, S.J. Therapies targeting DNA and RNA in Huntington’s disease. Lancet Neurol 16, 837–847 (2017).

6. Aylward, E.H., et al. Longitudinal change in basal ganglia volume in patients with Huntington’s disease. Neurology 48, 394–399 (1997).

7. Aylward, E.H. Change in MRI striatal volumes as a biomarker in preclinical Huntington’s disease. Brain Res Bull 72, 152–158 (2007).

8. Tabrizi, S.J., et al. Biological and clinical manifestations of Huntington’s disease in the longitudinal TRACK-HD study: cross-sectional analysis of baseline data. Lancet Neurol 8, 791–801 (2009).

9. Tabrizi, S.J., et al. Potential endpoints for clinical trials in premanifest and early Huntington’s disease in the TRACK-HD study: analysis of 24 month observational data. Lancet Neurol 11, 42–53 (2012).

10. Tabrizi, S.J., et al. Biological and clinical changes in premanifest and early stage Huntington’s disease in the TRACK-HD study: the 12-month longitudinal analysis. Lancet Neurol 10, 31–42 (2011).

11. Byrne, L.M., et al. Neurofilament light protein in blood as a potential biomarker of neurodegeneration in Huntington’s disease: a retrospective cohort analysis. Lancet Neurol 16, 601–609 (2017).

12. Byrne, L.M., et al. Cerebrospinal fluid neurogranin and TREM2 in Huntington’s disease. Sci Rep 8, 4260 (2018).

13. Byrne, L.M., et al. Evaluation of mutant huntingtin and neurofilament proteins as potential markers in Huntington’s disease. Sci Transl Med 10(2018).

14. Byrne, L.M. & Wild, E.J. Cerebrospinal Fluid Biomarkers for Huntington’s Disease. J Huntingtons Dis 5, 1–13 (2016).

15. Hua, J., Unschuld, P.G., Margolis, R.L., van Zijl, P.C. & Ross, C.A. Elevated arteriolar cerebral blood volume in prodromal Huntington’s disease. Mov Disord 29, 396–401 (2014).

16. Andrews, T.C., et al. Huntington’s disease progression. PET and clinical observations. Brain 122 (Pt 12), 2353–2363 (1999).

17. Ciarmiello, A., et al. Brain white-matter volume loss and glucose hypometabolism precede the clinical symptoms of Huntington’s disease. J Nucl Med 47, 215–222 (2006).

18. Feigin, A., et al. Thalamic metabolism and symptom onset in preclinical Huntington’s disease. Brain 130, 2858–2867 (2007).

19. Lopez-Mora, D.A., et al. Striatal hypometabolism in premanifest and manifest Huntington’s disease patients. Eur J Nucl Med Mol Imaging 43, 2183–2189 (2016).

20. Paulsen, J.S. Functional imaging in Huntington’s disease. Exp Neurol 216, 272–277 (2009).

21. Young, A.B., et al. PET scan investigations of Huntington’s disease: cerebral metabolic correlates of neurological features and functional decline. Ann Neurol 20, 296–303 (1986).

22. Weir, D.W., Sturrock, A. & Leavitt, B.R. Development of biomarkers for Huntington’s disease. Lancet Neurol 10, 573–590 (2011).

23. van den Bogaard, S., Dumas, E., van der Grond, J., van Buchem, M. & Roos, R. MRI biomarkers in Huntington’s disease. Front Biosci (Elite Ed) 4, 1910–1925 (2012).

24. Versluis, M.J., van der Grond, J., van Buchem, M.A., van Zijl, P. & Webb, A.G. High-field imaging of neurodegenerative diseases. Neuroimaging Clin N Am 22, 159–171, ix (2012).

25. Mergenthaler, P., Lindauer, U., Dienel, G.A. & Meisel, A. Sugar for the brain: the role of glucose in physiological and pathological brain function. Trends Neurosci 36, 587–597 (2013).

26. Gonzalez, R.G., et al. Functional MR in the evaluation of dementia: correlation of abnormal dynamic cerebral blood volume measurements with changes in cerebral metabolism on positron emission tomography with fludeoxyglucose F 18. AJNR Am J Neuroradiol 16, 1763–1770 (1995).

27. Lee, H., et al. Cell Type-Specific Transcriptomics Reveals that Mutant Huntingtin Leads to Mitochondrial RNA Release and Neuronal Innate Immune Activation. Neuron (2020).

28. Hua, J., et al. MRI techniques to measure arterial and venous cerebral blood volume. Neuroimage 187, 17–31 (2019).

29. Iadecola, C. & Nedergaard, M. Glial regulation of the cerebral microvasculature. Nat Neurosci 10, 1369–1376 (2007).

30. Kim, T., Hendrich, K.S., Masamoto, K. & Kim, S.G. Arterial versus total blood volume changes during neural activity-induced cerebral blood flow change: implication for BOLD fMRI. J Cereb Blood Flow Metab 27, 1235–1247 (2007).

31. Yang, S., et al. CRISPR/Cas9-mediated gene editing ameliorates neurotoxicity in mouse model of Huntington’s disease. J Clin Invest 127, 2719–2724 (2017).

32. Ekman, F.K., et al. CRISPR-Cas9-Mediated Genome Editing Increases Lifespan and Improves Motor Deficits in a Huntington’s Disease Mouse Model. Mol Ther Nucleic Acids 17, 829–839 (2019).

33. Menalled, L. & Brunner, D. Animal models of Huntington’s disease for translation to the clinic: best practices. Mov Disord 29, 1375–1390 (2014).

34. Menalled, L.B. Knock-in mouse models of Huntington’s disease. NeuroRx 2, 465–470 (2005).

35. Gray, M. Astrocytes in Huntington’s Disease. Adv Exp Med Biol 1175, 355–381 (2019).

36. Drouin-Ouellet, J., et al. Cerebrovascular and blood-brain barrier impairments in Huntington’s disease: Potential implications for its pathophysiology. Ann Neurol 78, 160–177 (2015).

37. Rieux, M., et al. Shedding a new light on Huntington’s disease: how blood can both propagate and ameliorate disease pathology. Mol Psychiatry (2020).

38. Menalled, L.B., et al. Comprehensive behavioral and molecular characterization of a new knock-in mouse model of Huntington’s disease: zQ175. PLoS One 7, e49838 (2012).

39. Heikkinen, T., et al. Characterization of neurophysiological and behavioral changes, MRI brain volumetry and 1H MRS in zQ175 knock-in mouse model of Huntington’s disease. PLoS One 7, e50717 (2012).

40. Peng, Q., et al. Characterization of Behavioral, Neuropathological, Brain Metabolic and Key Molecular Changes in zQ175 Knock-In Mouse Model of Huntington’s Disease. PLoS One 11, e0148839 (2016).

41. Vanlandewijck, M., et al. A molecular atlas of cell types and zonation in the brain vasculature. Nature 554, 475–480 (2018).

42. Bankhead, P., Scholfield, C.N., McGeown, J.G. & Curtis, T.M. Fast retinal vessel detection and measurement using wavelets and edge location refinement. PLoS One 7, e32435 (2012).

43. Bjornsson, C.S., et al. Associative image analysis: a method for automated quantification of 3D multi-parameter images of brain tissue. J Neurosci Methods 170, 165–178 (2008).

44. Kerschnitzki, M., et al. Architecture of the osteocyte network correlates with bone material quality. J Bone Miner Res 28, 1837–1845 (2013).

45. Lin, C.Y., et al. Neurovascular abnormalities in humans and mice with Huntington’s disease. Exp Neurol 250, 20–30 (2013).

46. Zacchigna, S., Lambrechts, D. & Carmeliet, P. Neurovascular signalling defects in neurodegeneration. Nat Rev Neurosci 9, 169–181 (2008).

47. Cepeda-Prado, E., et al. R6/2 Huntington’s disease mice develop early and progressive abnormal brain metabolism and seizures. J Neurosci 32, 6456–6467 (2012).

48. Hua, J., et al. Inflow-based vascular-space-occupancy (iVASO) MRI. Magn Reson Med 66, 40-56 (2011).

49. Niccolini, F. & Politis, M. Neuroimaging in Huntington’s disease. World J Radiol 6, 301–312 (2014).

50. Pouladi, M.A., Morton, A.J. & Hayden, M.R. Choosing an animal model for the study of Huntington’s disease. Nat Rev Neurosci 14, 708–721 (2013).

51. Parievsky, A., Cepeda, C. & Levine, M.S. Evidence from the R6/2 Mouse Model of Huntington’s Disease for Using Abnormal Brain Metabolism as a Biomarker for Evaluating Therapeutic Approaches for Treatment. Future Neurol 7, 527–530 (2012).

52. Li, X., et al. Association of Glioma Grading With Inflow-Based Vascular-Space-Occupancy MRI: A Preliminary Study at 3T. J Magn Reson Imaging 50, 1817–1823 (2019).

53. Paez, A., et al. Differential changes in arteriolar cerebral blood volume between Parkinson’s disease patients with normal and impaired cognition and individuals with mild cognitive impaired (MCI) due to Alzheimer’s disease– an exploratory study. Tomography (2020).

54. Hua, J., et al. Increased cerebral blood volume in small arterial vessels is a correlate of amyloid-beta-related cognitive decline. Neurobiol Aging 76, 181–193 (2019).

55. Hua, J., et al. Abnormal Grey Matter Arteriolar Cerebral Blood Volume in Schizophrenia Measured With 3D Inflow-Based Vascular-Space-Occupancy MRI at 7T. Schizophr Bull 43, 620–632 (2017).

56. Iturria-Medina, Y., et al. Early role of vascular dysregulation on late-onset Alzheimer’s disease based on multifactorial data-driven analysis. Nat Commun 7, 11934 (2016).

57. Lu, H., Donahue, M.J. & van Zijl, P.C. Detrimental effects of BOLD signal in arterial spin labeling fMRI at high field strength. Magn Reson Med 56, 546–552 (2006).

58. Poplawsky, A.J., et al. Dominance of layer-specific microvessel dilation in contrast-enhanced high-resolution fMRI: Comparison between hemodynamic spread and vascular architecture with CLARITY. Neuroimage 197, 657–667 (2019).

59. Walther, N., et al. A quantitative map of human Condensins provides new insights into mitotic chromosome architecture. J Cell Biol 217, 2309–2328 (2018).

