## Supplemental figures 1-3 for "*HTT* silencing delays onset and slows progression of Huntington’s disease like phenotype: Monitoring with a novel neurovascular biomarker"

### Extended Figure 1

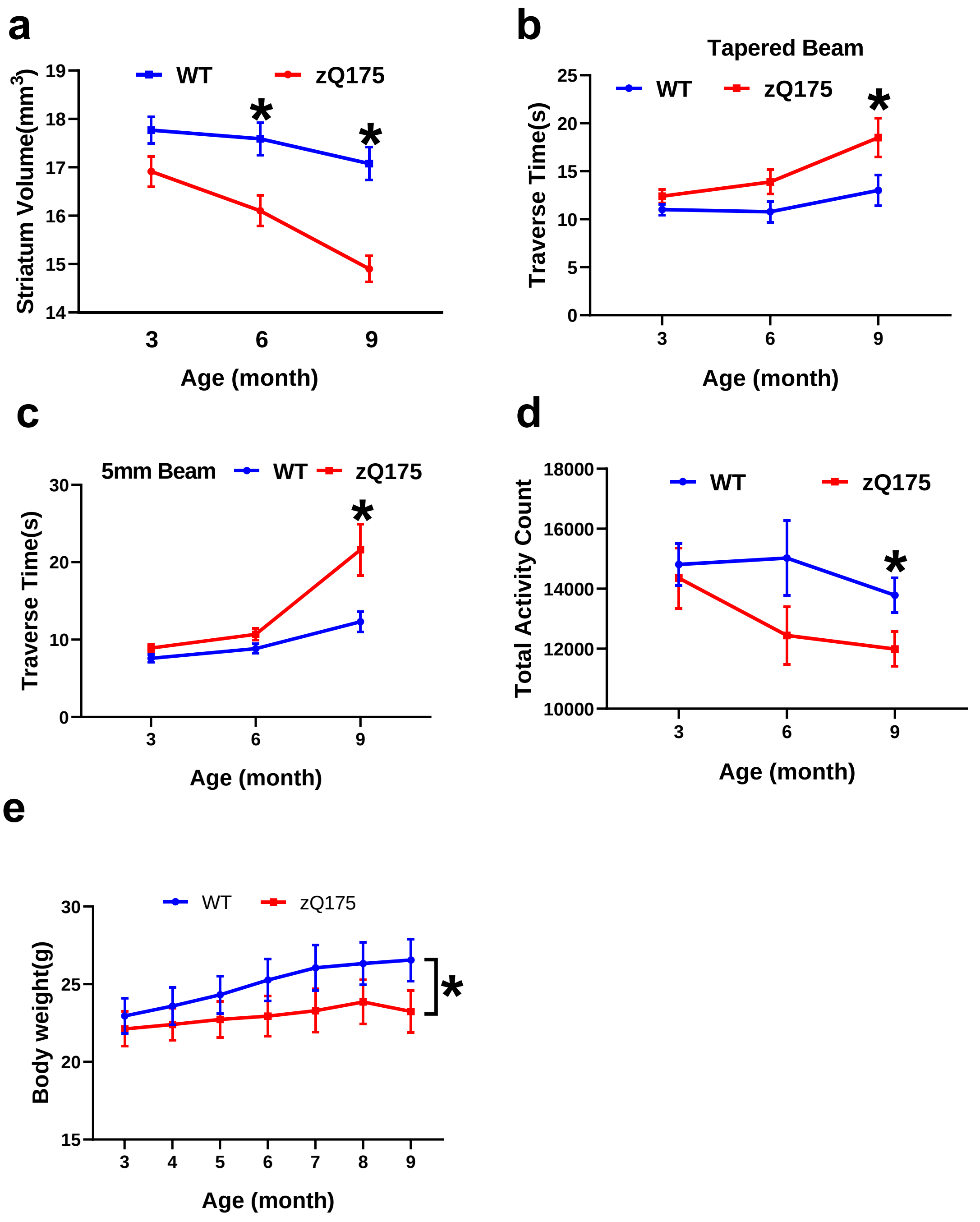

### Extended Figure 2

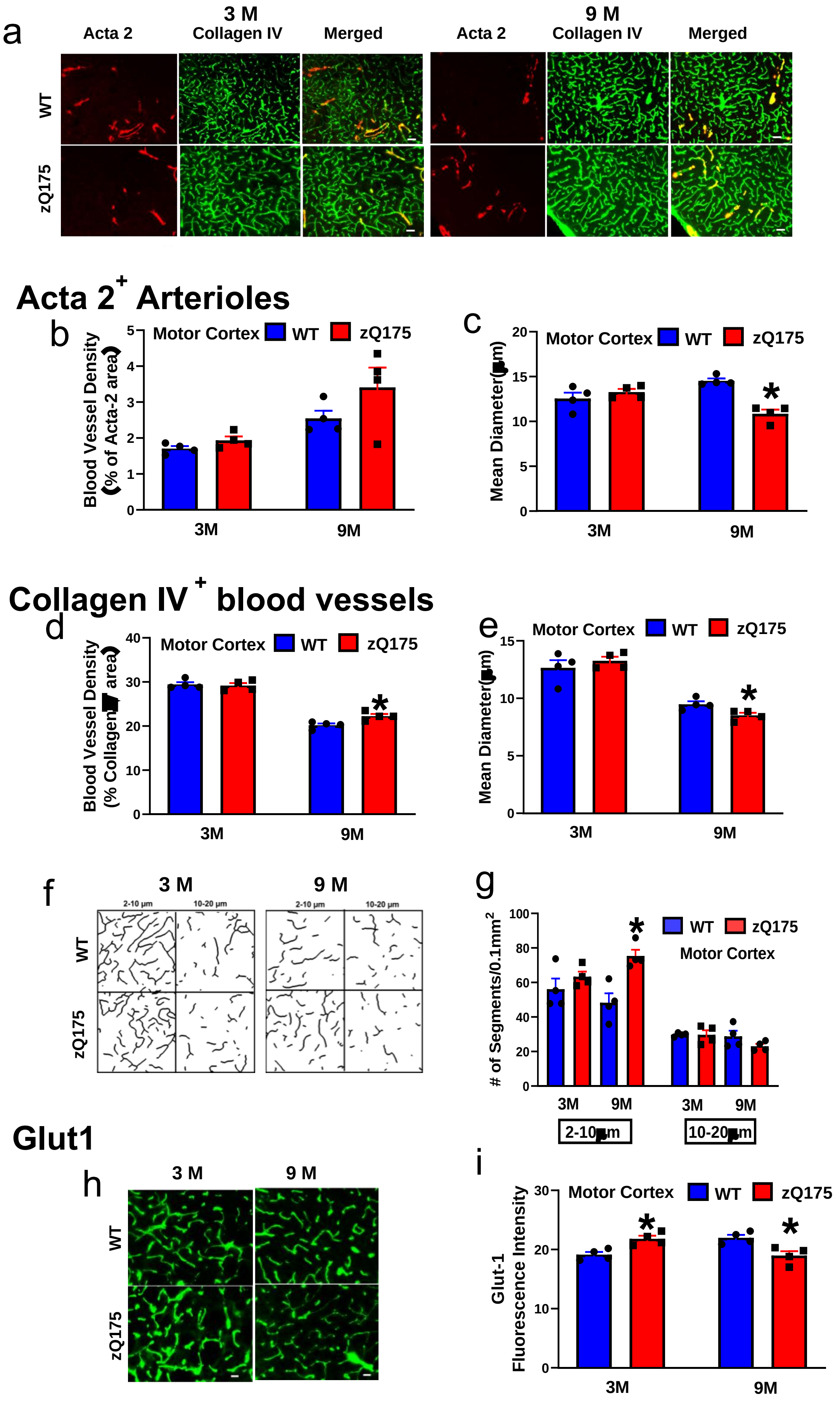

### Extended Figure 3

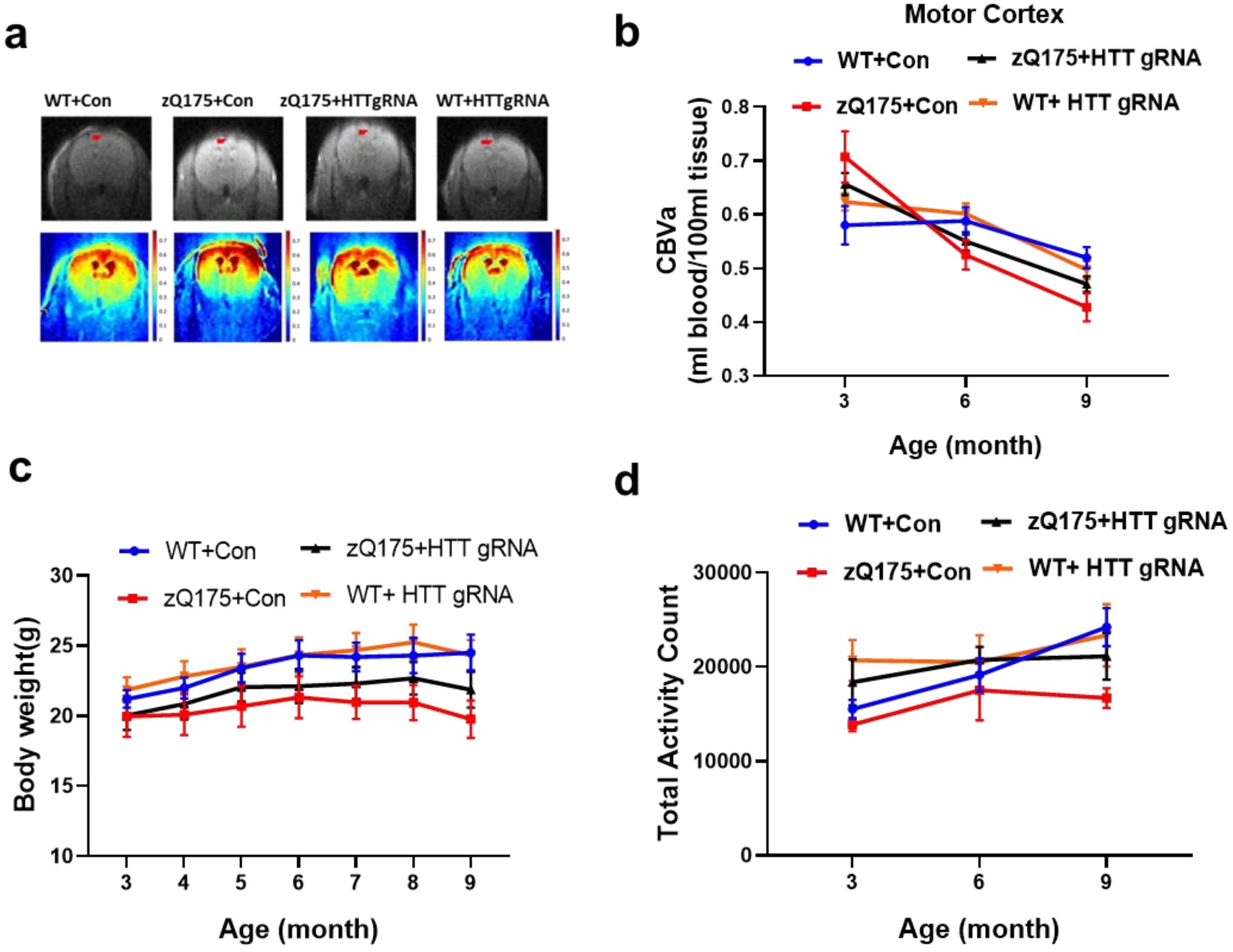

**Extended data Figure 1. Longitudinal characterization of heterozygous zQ175 mice. (a)** The volume of the striatum was measured by structural MRI in the mice at indicated ages and genotype. (**b)** Mice were tested on a tapered beam and time crossing the beam (Traverse time) was recorded. (**c**) Mice were tested on a 5 mm balance beam and time crossing the beam (Traverse time) was recorded. (**d**) Locomotor activity was assessed in an open field apparatus. Total activity was recorded automatically during a 1 h testing period. (**e**) Body weight was recorded monthly. All data are Mean ± SEM, **p* < 0.05 *versus* the values of WT group at the corresponding ages by the one-way (genotype) *ANOVA* or longitudinal fata with Two-way (age and genotype) *ANOVA* with Bonferroni *post hoc* analysis.

**Extended data Figure 2. Morphological analysis of cerebral vasculature in the motor cortex of zQ175 mice. (a)** Representative images of immunostaining signals of Acta2 (Red, arterioles) and collagen IV (Green, all blood vessels) in the mouse motor cortex from indicated genotypes and ages. 3 months (3M) represents premanifest stage of zQ175 mice; 9 months (9M) represents manifest zQ175 mice. Scale bar = 20 µm. (**b-c**) Quantitative analysis of Acta $2^+$ arteriolar density (b) and mean diameter (c) in the motor cortex of zQ175 mice and wild type (WT) controls at 3 and 9 months of age. (**d-e**) Quantitative analysis of collagen $IV^+$ blood vessel density (d) and mean diameter (e). (**f**) The representative skeletonized image of collagen $IV^+$ blood vessels with indicated diameters in the zQ175 and WT mice at 3 and 9 months of age. (**g**) Quantitative analysis of the number of segments per 0.1 $mm^2$ in collagen $IV^+$ blood vessels at indicated diameters. (**h-i**) Representative images of Glut1 immunostaining in the motor cortex of mice from indicated genotypes and ages (h) and quantitative data of Glut1 immunofluorescent intensity (i). Scale bar = 20 µm. All data are Mean ± SEM, n = 4 mice/group. **p*<0.05 *versus* the values of WT group at the corresponding ages by Student's *t*-tests.

**Extended data Figure 3**. **Effect of CRISPR/Cas9-mediated HTT silencing in the striatal neurons on CBVa in the motor cortex, body weight and locomotor activity in zQ175 mice**. (**a**)

Representative CBVa maps calculated from iVASO images in the motor cortex of mice from indicated groups. Top row shows the raw images, and the red ROIs indicate the quantified motor cortex region. Bottom row shows the CBVa maps for indicated groups at 3 months of age. The scale bars are shown on the right and warmer color represents higher CBVa values. (**b**) Quantification of CBVa values in the motor cortex from indicated groups and ages. (**c**) Longitudinal body weight in mice from indicated genotype and treatment. (**d**) Longitudinal locomotor activity data in an open field apparatus. All data are Mean ± SEM, n = 6-9 mice/group.
